# The bladder cancer m^6^A landscape is defined by methylation dilution and 3′-UTR hypermethylation

**DOI:** 10.1101/2025.07.11.664345

**Authors:** Jonas Koch, Jinyun Xu, Felix Bormann, Vitor Coutinho Carneiro, Manuel Neuberger, Katja Nitschke, Malin Nientiedt, Philipp Erben, Maurice Stephan Michel, Manuel Rodriguez-Paredes, Frank Lyko

## Abstract

N^6^-Methyladenosine (m^6^A) is the most abundant internal modification of eukaryotic mRNAs and regulates target transcripts throughout the mRNA life cycle. Although changes in m^6^A have been reported in human cancers, technical limitations have hindered a comprehensive understanding of the cancer-associated m^6^A landscape. Here, we used GLORI-sequencing to establish the first transcriptome-wide, single-nucleotide resolution maps of m^6^A in cancer. Differentially methylated transcripts were enriched in oncogenic pathways relevant to UCB. We discovered two key m^6^A signatures in UCB: a global loss of methylation and a local hypermethylation at 3′-UTRs. Integration of RNA-sequencing data revealed that the global loss resulted from dilution of methylation marks due to increased expression of unmethylated transcripts and decreased expression of highly methylated transcripts. In contrast, local 3′-UTR hypermethylation was associated with the overexpression of VIRMA, a component of the m^6^A writer complex, which was linked to UCB progression. Our study is the first to describe the m^6^A epitranscriptomic landscape of cancer at single-base resolution and provides first insights into the processes that generate its characteristic signatures.

## Introduction

N^6^-Methyladenosine (m^6^A) is the most frequent internal modification of mammalian mRNAs and is present in 0.15-0.6 % of all adenosine residues (He & He, 2021). The modification is deposited in the nucleus by the m^6^A writer complex, which consists of the METTL3, METTL14, WTAP, VIRMA, ZC3H13, CBLL1, RBM15, and RBM15B proteins (Zaccara *et al*, 2019). METTL3 represents the catalytically active writer enzyme, with the other components of the complex being responsible for RNA binding and additional regulatory functions (Zaccara *et al*., 2019). While the mechanisms underlying m^6^A patterning across the transcriptome remain poorly understood, results from HeLa cells have suggested that VIRMA acts as a regulatory subunit that recruits the m^6^A writer complex to 3-′UTRs and stop codon regions (Yue *et al*, 2018). m^6^A is predominantly found in the DRAC(H) consensus sequence, in which D = A, G or T; R = A or G; and H = A, C or U (Dominissini *et al*, 2012; Linder *et al*, 2015). The modification is read by the proteins of the YTH family, which comprises YTHDC1, YTHDF1, YTHDF2, and YTHDF3. These proteins were also shown to mediate m^6^A-dependent downstream functions (Flamand *et al*, 2023; Zaccara *et al*., 2019). In this context, different molecular mechanisms of m^6^A-dependent transcript regulation were described, including alternative splicing, alternative polyadenylation, transport, stability, and translation (Boulias & Greer, 2023; Flamand *et al*., 2023; Koch & Lyko, 2024). Importantly, deregulation of m^6^A has been found to affect physiological and pathophysiological processes, including cancer development (Lan *et al*, 2019).

Bladder cancer is a major global health problem and the 10^th^ most frequent cancer worldwide (Saginala *et al*, 2020). As 90-95 % of bladder cancers originate from urothelial cells, urothelial carcinoma of the bladder (UCB) represents the most common form of bladder cancer (Dyrskjøt *et al*, 2023; Siegel *et al*, 2024). About 75 % of all patients are diagnosed with non-muscle-invasive bladder cancer (NMIBC). 10-20 % of NMIBC cases further progress to muscle-invasive bladder cancer (MIBC), which is characterized by low survival rates and a high metastatic potential (Berdik, 2017; Lopez-Beltran *et al*, 2024; Sylvester *et al*, 2006). Approved early detection biomarkers for UCB are currently not available (Batista *et al*, 2020; Gill & Perks, 2024). Also, UCB incidence and prevalence numbers are expected to increase due to population growth and aging (Richters *et al*, 2020). Therefore, novel therapeutic drug targets and biomarkers are urgently needed for improving patient prognosis.

Multiple studies reported evidence for cancer-associated changes in the m^6^A epitranscriptome (Deng *et al*, 2022; Deng *et al*, 2023). This includes aberrant expression of m^6^A regulators and changes in the m^6^A methylation of individual transcripts related to UCB development (Yang *et al*, 2024b). Using LC-MS/MS analysis, we have shown recently that m^6^A levels were strongly reduced in UCB when compared to matched paratumoral tissue (Koch *et al*, 2023). This presents an apparent contradiction to other published findings where local hypermethylation of cancer-associated transcripts was described (Cheng *et al*, 2019). To resolve this inconsistency and to provide detailed maps of m^6^A in cancer, novel and robust m⁶A detection techniques are required, as the previous studies relied on antibody-dependent mapping techniques with limited resolution, lack of quantitative information, and insufficient specificity (Helm *et al*, 2019; Koch & Lyko, 2024; McIntyre *et al*, 2020). Similarly, initial attempts to map m^6^A distribution by direct RNA-sequencing were limited by low sequencing coverage and robustness (Li *et al*, 2024; Zhang *et al*, 2024). These methodological limitations have greatly limited our understanding of cancer-associated changes in m^6^A patterning and their functional implications in cancer development and progression.

GLORI-sequencing is a newly established method that allows for the site-specific, absolute quantification of m^6^A through an unbiased chemical deamination protocol (Liu *et al*, 2023). As similar methods, such as whole-genome bisulfite sequencing (Lister & Ecker, 2009) have been highly successful in mapping the cancer epigenome, we adopted GLORI to generate the first transcriptome-wide, base-resolution maps of m^6^A in cancer. We sequenced and compared a set of N=18 clinical UCB samples and non-malignant uroepithelial control samples and uncovered systematic differences in the m^6^A landscape of UCB. More specifically, the integration of RNA expression and m^6^A methylation data revealed a global dilution of methylation in UCB tissues. Simultaneously, UCB samples showed local 3’-UTR hypermethylation, which correlated with alternative mRNA polyadenylation. Interestingly, these changes were associated with the overexpression of VIRMA, a component of the m^6^A writer complex, which we found to be clinically relevant for the progression of UCB. Our findings thus uncover key signatures of the cancer m^6^A epitranscriptome and provide first insight into the underlying mechanisms.

## Results

### Establishment of a m^6^A methylation analysis pipeline

In an initial set of experiments, we attempted to reproduce the published GLORI results from HEK293T cells (Liu *et al*., 2023). Because the guanosine protection step leads to substantial RNA degradation (Sun *et al*, 2025), we adjusted the original protocol to improve library quality (see Methods for details). Subsequently, we sequenced libraries from three replicates, with an average yield of 200 million sequencing reads, respectively, and mapping rates of 50 % (Table EV 1). Importantly, we obtained median A-to-G conversion ratios of >99 % (Table EV 2), which is similar to the reported conversion ratios and strongly limits the influence of false-positive signals (Liu *et al*., 2023). In total, we detected a shared 69,748 m^6^A sites in the transcriptomes of the three HEK293T cell replicates (Figure EV 1A). The majority (87 %) of these sites were also detected previously (Figure EV 1B), with differences being largely due to the higher sequencing depth of the original study (Liu *et al*., 2023). Also, methylation levels were found to be highly reproducible (Figure EV 1C). Finally, it should be noted that our results are in excellent agreement with results that were obtained by direct RNA-sequencing of the same batch of cells (Hewel *et al*, 2025), which provides important orthogonal validation for our GLORI-based mapping and quantification of m^6^A sites.

In the next step, we repeated the protocol for the T24 UCB cell line and again obtained high-quality GLORI datasets (Table EV 1 and Table EV 2). Data analysis identified more than 88,000 m^6^A sites, which were predominantly found in the DRAC(H) consensus sequence motif (Fig. 1A). Within the 15 most frequent pentanucleotide motifs, canonical DRAC(H) motifs were strongly overrepresented (Fig. 1B). Also, higher and more evenly distributed m^6^A methylation levels were detected in the DRAC(H) motifs, while the non-canonical motifs were lowly methylated (Fig. 1C). Based on these observations, we considered the m^6^A sites in DRAC motifs as high-confidence methylation marks, while the m^6^A sites detected in non-canonical motifs were interpreted as deamination or sequencing artefacts. We therefore restricted all further analyses to m^6^A sites detected in DRAC consensus sequence motifs. Calculating the number of m⁶A sites per transcript showed a median of n=4, but several transcripts were found to have multiple (up to 154) m^6^A sites (Fig. 1D). Overall, DRAC motifs showed a bimodal methylation distribution with peaks at 20 % and 95 % methylation (Fig. 1E). Metagene plots showed that the majority of m^6^A sites were in the CDS and 3’-UTR of the transcripts with a characteristic peak around the stop codon (Fig. 1F). Treatment of T24 cells with the METTL3 inhibitor STM2457 resulted in strongly reduced methylation, both transcriptome-wide (Fig. 1G) and at the level of specific transcripts (Fig. 1H). These findings demonstrate the capacity of GLORI to faithfully map m^6^A sites.

**Figure 1.**
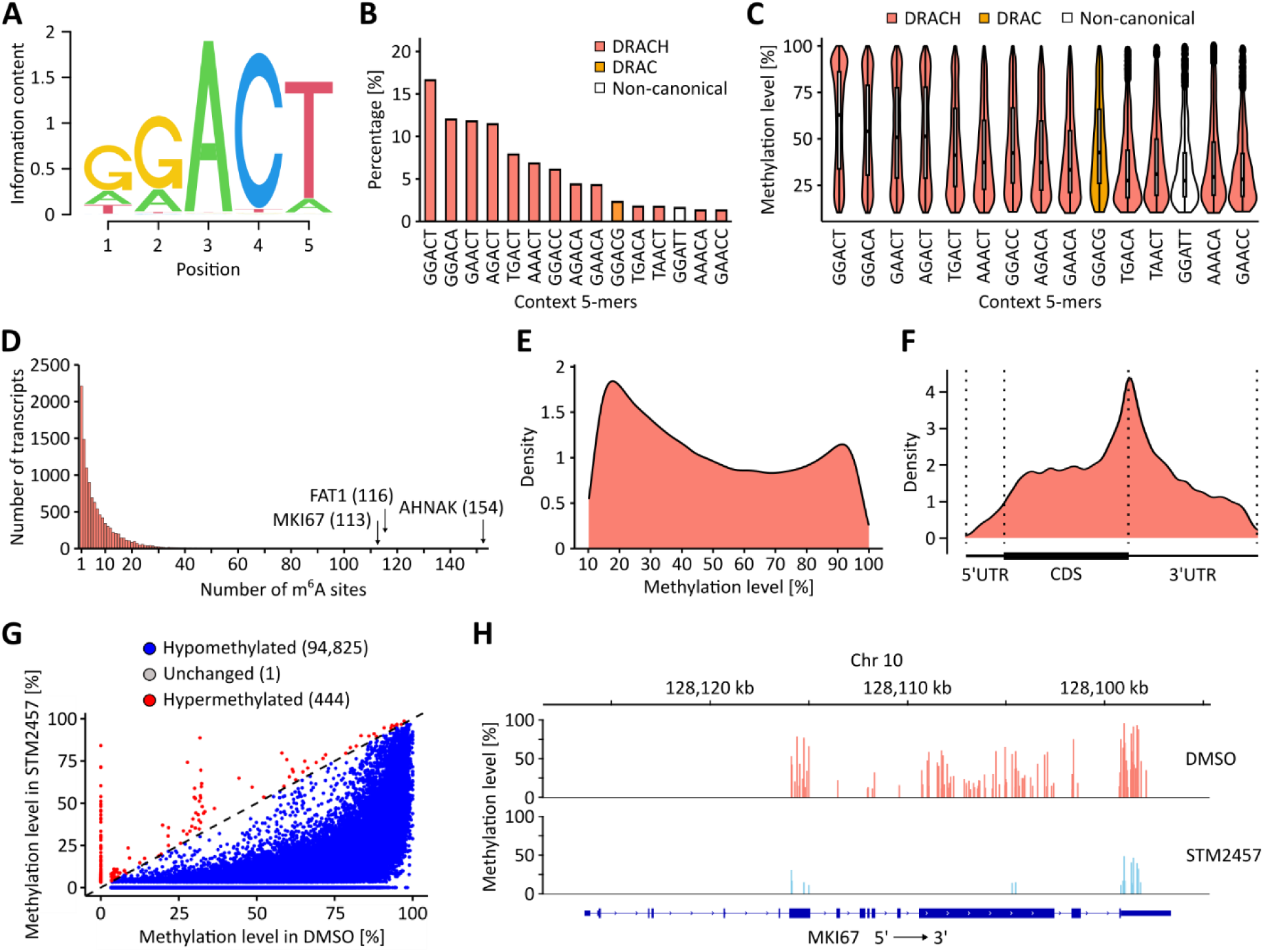
GLORI-based analysis of the m^6^A landscape in T24 UCB cells. **(A)** Sequence motif analysis of m^6^A sites revealing the DRAC(H) consensus motif. **(B)** Frequency of the 15 most detected m^6^A site motifs. DRAC(H) motifs were detected more frequently than non-canonical motifs. **(C)** Quantification of the m^6^A methylation level in the 15 most detected m^6^A site motifs. The lowest m^6^A methylation levels were detected non-canonical motifs. **(D)** Quantification of m^6^A sites per transcript. While the median number was n=4, some transcripts harbored very high numbers of m^6^A sites, as indicated. **(E)** Kernel density plot showing that the majority of m^6^A sites have either low or high methylation levels. **(F)** Metaplot showing that detected m^6^A sites predominantly occur in the CDS and 3’-UTR regions with a peak density surrounding the stop codon. **(G)** Scatter plot showing that METTL3 inhibitor STM2457 treatment resulted in reduced methylation levels for 94,825 sites, while 444 sites were found to have increased methylation levels. **(H)** IGV browser tracks showing the detected m^6^A sites in the MKI67 transcript. STM2457 treatment led to a pronounced reduction of m^6^A methylation.

### Comparative analysis of UCB and control samples

In the next step, we applied our GLORI pipeline to nine UCB and nine non-malignant uroepithelial samples (control). Clinical information about the patient samples is provided in Table EV 3. The quality of the corresponding GLORI datasets matched the high standards obtained with the HEK293T and T24 cell lines (Table EV 1 and Table EV 2). Initial sequence motif analyses confirmed methylation in the consensus DRAC(H) motif for both sample groups (Fig. 2A) and did not reveal any detectable differences in pentanucleotide motif frequencies and methylation levels (Fig. 2B and Fig. 2C). These results strongly suggest that m^6^A motif specificity is retained in UCB. In total, approximately 42,000 detected m^6^A sites were shared among the UCB samples, while roughly 45,000 m^6^A sites were shared among the control samples and used for further downstream analysis. When comparing the distribution of methylation levels (Fig. 2D) and the localization of m^6^A sites (Fig. 2E), we could not identify any major differences between UCB and control samples. Interestingly, however, principal component analysis based on all m^6^A sites showed that the tumor samples could be clearly separated from the controls (Fig. 2F and Figure EV 2A). These findings strongly suggest the presence of cancer-specific signatures in the m^6^A epitranscriptomic landscape.

**Figure 2.**
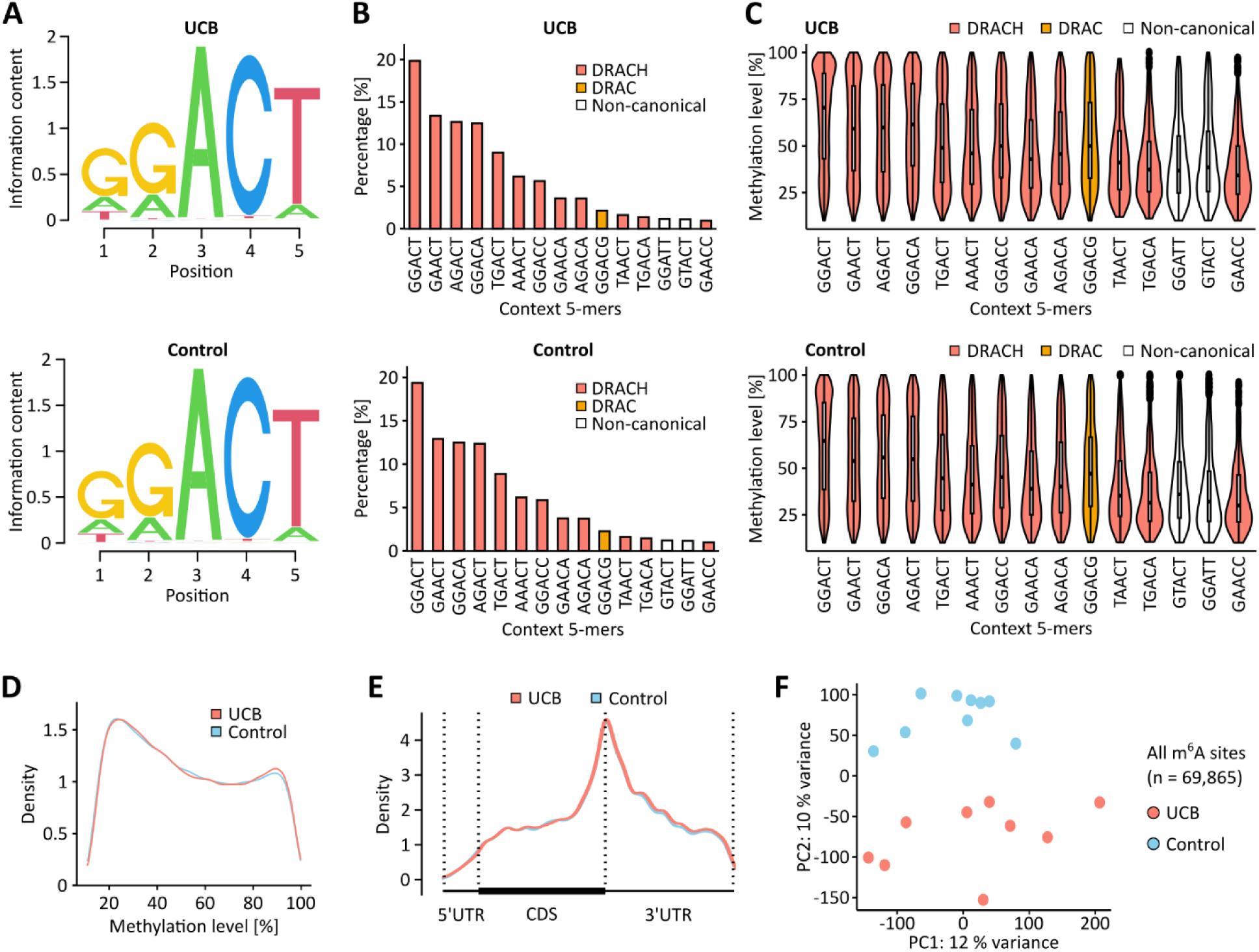
The m^6^A epitranscriptomic landscapes of UCB and non-malignant uroepithelial tissue show moderate, but systematic differences. **(A)** Sequence motif analyses using detected m^6^A sites in UCB and control tissue samples. In both tissue types, m^6^A sites were primarily detected in the DRAC(H) consensus sequence motif. **(B)** Frequencies of the 15 most detected motifs. **(C)** Quantification of the methylation level in the 15 most detected motifs. **(D)** Kernel density plot showing the distribution of m^6^A site methylation levels in the UCB and control data sets. **(E)** Metagene plots demonstrating that the detected m^6^A sites predominantly occur in CDS and 3’-UTR regions independent of the tissue origin. **(F)** Principal component analysis demonstrating that UCB and control tissue samples can be separated based on their m^6^A signatures.

### Differential methylation of cancer-related transcripts

In the next step, we sought to further investigate the cancer-associated m^6^A pattern changes by identifying differentially methylated m^6^A sites using stringent criteria (|Δm^6^A|≥10 % and p<0.05). This identified 1,921 sites with increased methylation levels in cancer (hypermethylated) and 1,238 sites with reduced methylation levels in cancer (hypomethylated, Fig. 3A). Principal component and additional dimensionality reduction analyses based on these 3,195 differentially methylated sites again showed a clear separation of the tumor and control sample groups (Fig. 3B and Figure EV 2B). On the transcript level, 1,186 transcripts were identified to be hypermethylated, while 902 transcripts were hypomethylated (Fig. 3C). Pathway analyses showed that the differentially methylated transcripts were enriched in several pathways that have been linked to UCB development (Goriki *et al*, 2018; Knowles & Hurst, 2015; Sui *et al*, 2017), including TNFα, NOTCH, p53, and TGFβ signaling, as well as pathways related to apoptosis and epithelial mesenchymal transition (EMT, Fig. 3D). Additional differentially methylated transcripts included MYC, SMAD3, and BTG2, which have all been linked to UCB (Cheng *et al*., 2019; Mao *et al*, 2015; Millet & Zhang, 2007; Yuniati *et al*, 2019) and were found hypermethylated in their CDS and 3’-UTR regions (Fig. 3E and Figure EV 3).

**Figure 3.**
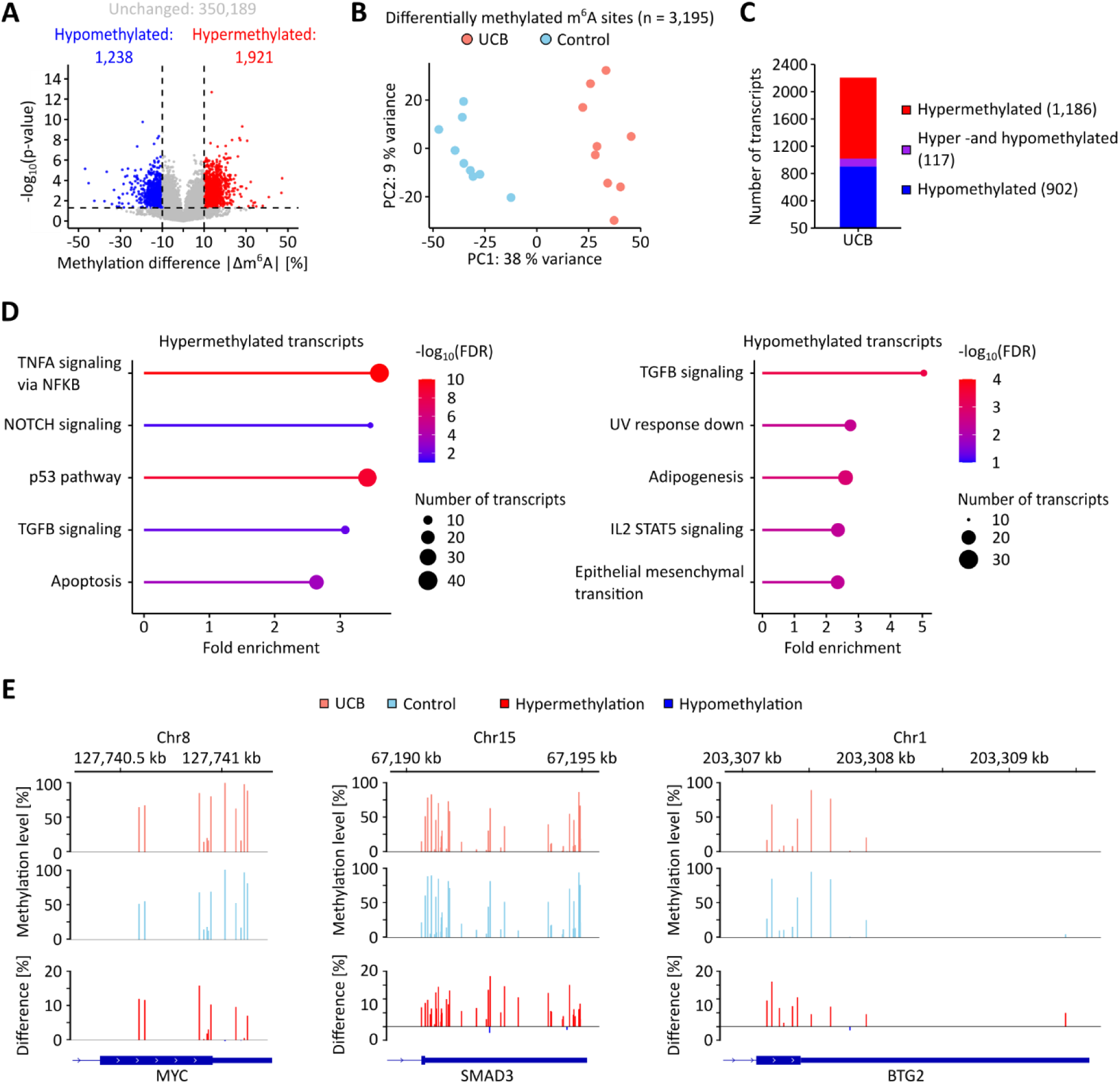
The UCB m^6^A epitranscriptome is characterized by both hypo- and hypermethylation. **(A)** Differential methylation analysis revealed that 1,921 m^6^A sites were hypermethylated, while 1,238 m^6^A sites were hypomethylated. Thresholds: Absolute difference in methylation level >10 %, p<0.05. **(B)** PCA showing that patient samples can be separated based on the differentially methylated m^6^A sites. **(C)** On the transcript level, 1,186 transcripts were found to be hypermethylated, 117 transcripts had both hyper - and hypomethylated m^6^A sites, and 902 transcripts were hypomethylated. **(D)** Top 5 enriched pathways based on differentially methylated transcripts. **(E)** Prominent examples for hypermethylated transcripts in UCB.

### Cancer-associated hypomethylation is due to cancer-associated changes in transcript abundance

As we observed both hypomethylation and hypermethylation in cancer samples, we performed further analyses to define the signatures of the cancer-associated m^6^A epitranscriptome. To address cancer-related differences in transcript abundance, we performed RNA-sequencing on the same tissue samples that we had used for GLORI. Principal component analysis showed that both sample groups could be separated based on their gene expression profiles (Figure EV 4A). Differential gene expression analysis demonstrated pronounced global deregulation of gene expression in UCB when compared to the control tissue, with 9,208 differentially expressed genes (Figure EV 4B). Cumulative analysis of normalized transcript levels showed that few transcripts made up a high proportion of the total transcriptome (Fig. 4A), with roughly 100 transcripts making up 50 % of the transcriptome in both groups. This raised the possibility that methylation and/or expression changes in these highly abundant transcripts could strongly impact the global methylation level. Indeed, integrated analysis of GLORI- and RNA-sequencing data showed a significant (p<0.05, t-test) global hypomethylation of UCB tissue samples (Fig. 4B), which confirmed the results observed in our previous LC-MS/MS analyses (Koch *et al*., 2023). Further data analysis revealed that unmethylated, highly abundant transcripts were upregulated, while highly methylated, highly abundant transcripts were downregulated in UCB (Fig. 4C). For example. several highly methylated transcripts (aggregate methylation >2), including EGR1, JUN, JUNB, and FOS, were markedly downregulated in UCB (Fig. 4D). In contrast, several unmethylated transcripts from the S100 family were upregulated in UCB (Fig. 4E) These findings strongly suggest that m^6^A marks become “diluted” in cancer samples, due to the increased presence of unmethylated transcripts.

**Figure 4.**
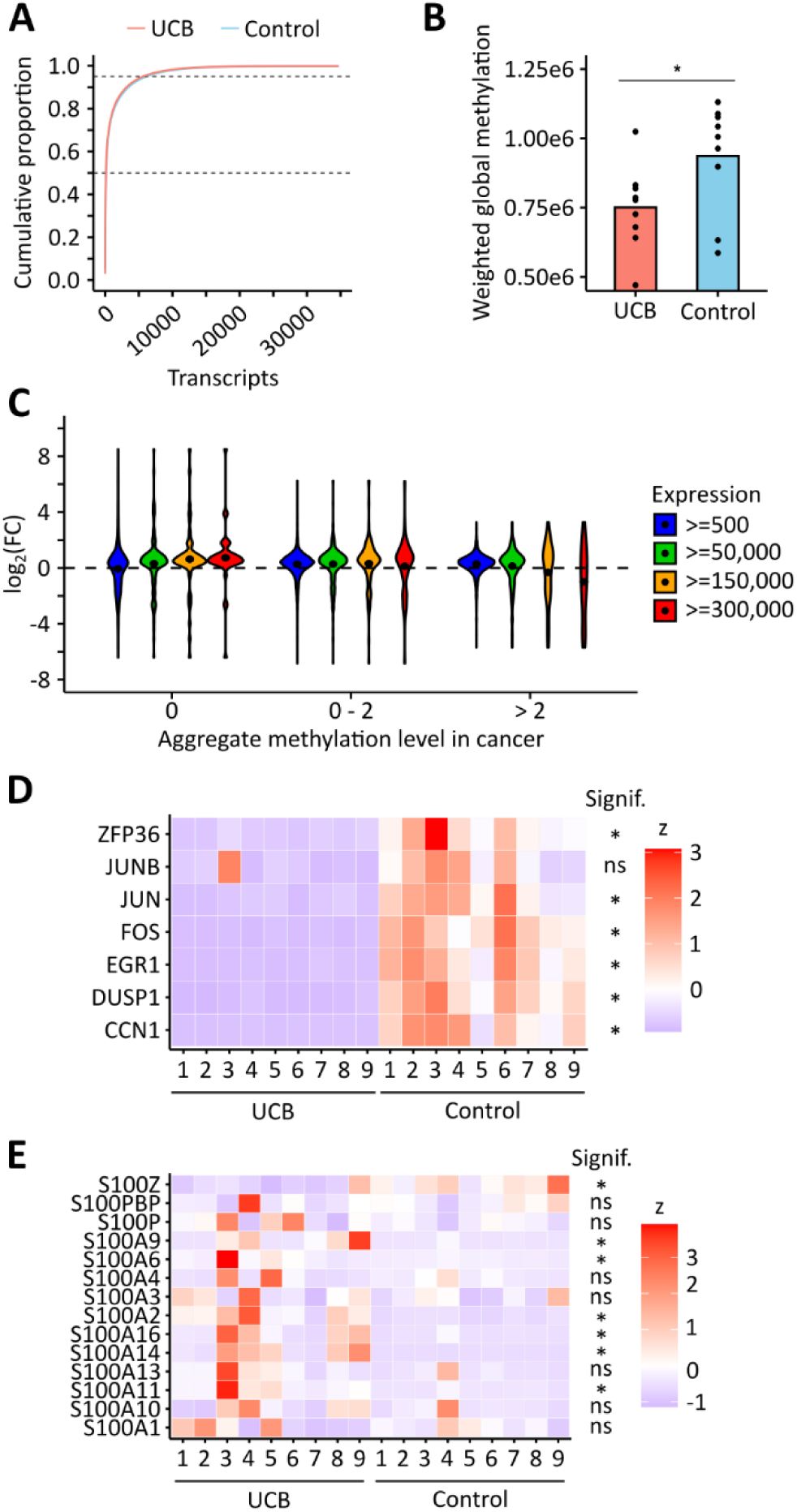
Global m^6^A hypomethylation in UCB results from changes in transcript abundance. **(A)** Cumulative proportion plot of control and UCB tissues. Few transcripts make up a high proportion of the transcriptome in both conditions. **(B)** Weighted global methylation levels of control and UCB tissues. The analysis confirms a global hypomethylation in UCB. p<0.05, t-test. **(C)** Global analysis of transcript expression changes in UCB considering the methylation status of the transcripts. When analyzing all transcripts, no major trends were observed. When restricting the analysis to highly abundant transcripts, an upregulation of unmethylated transcripts as well as a downregulation of highly methylated transcripts was observed in UCB. **(D)** Heatmap showing selected transcripts that had highest methylation levels in the lists of the most abundant transcripts in control and UCB tissues. Six out of the seven transcripts were downregulated in UCB. Threshold: q<0.05. **(E)** Heatmap showing the expression of the S100 transcripts family. Several transcripts were found to be upregulated in UCB. Threshold: q<0.05.

### Cancer-associated hypermethylation is enriched at 3’-UTRs and associated with increased VIRMA expression

To further characterize the contrasting signature, i.e. cancer-associated hypermethylation, we performed sequence motif analyses. The results showed that hypermethylation was frequently detected in the GGACT motif, while hypomethylated m^6^A sites were often found in the TGACT and AAACT motifs (Fig. 5A). Furthermore, metagene analyses found that hypermethylated m^6^A sites were predominantly located in the 3’-UTR, especially in proximity to the stop codon, while no clear patterns were detectable for hypomethylated sites (Fig. 5B). As m^6^A methylation changes have been shown to affect mRNA polyadenylation (Yue *et al*., 2018), we analyzed our datasets for evidence of alternative polyadenylation. Indeed, the majority of transcripts had negative difference of Percentage of Distal polyA site Usage Indices (PDUIs) indicating that they were shortened (Fig. 5C). To test whether m^6^A could affect alternative polyadenylation in UCB, we expanded our alternative polyadenylation analyses to UCB METTL3 knockout cell clones and to UCB cell lines treated with the METTL3 inhibitor STM2457. RNA-sequencing data analysis showed that most transcripts had positive ΔPDUIs upon METTL3 knockout and inhibition, which corresponds to transcript lengthening (Fig. 5D and Figure EV 5). These results further suggest that changes in m^6^A methylation affect alternative polyadenylation. Of note, almost all transcripts with negative ΔPDUI were m^6^A-methylated in UCB (Fig. 5E). Taken together, these findings establish hypermethylation near stop codons as a second important signature of the cancer-associated m^6^A epitranscriptome.

**Figure 5.**
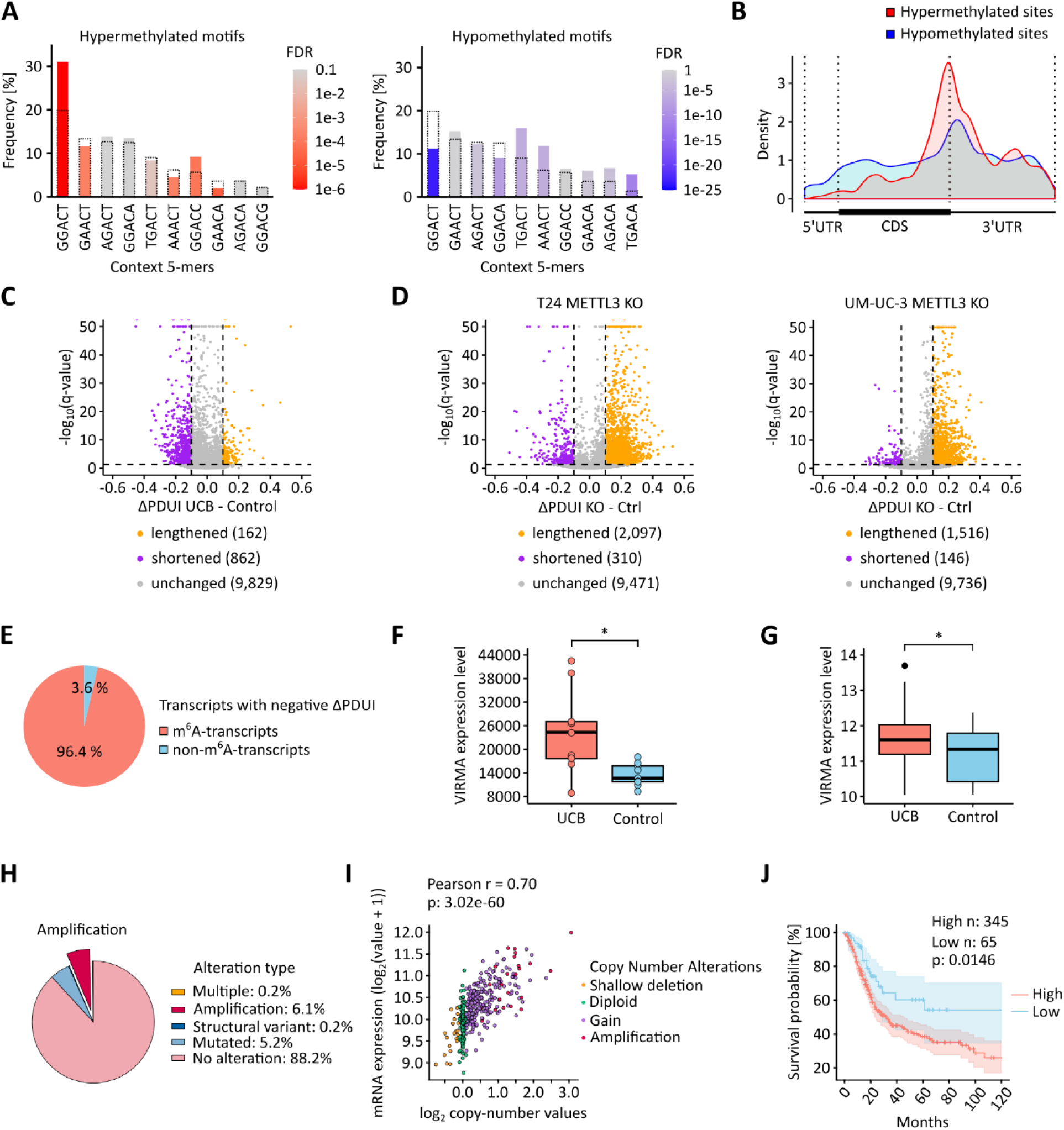
Local 3’-UTR hypermethylation is associated with an upregulation of VIRMA in UCB. **(A)** Frequency of the motifs detected in hypermethylated and hypomethylated m^6^A sites, respectively. Bars with dashed outlines represent the overall frequency of the respective motif in the cancer samples. Colored bars represent the frequency of the respective motif among the differentially methylated m^6^A sites. **(B)** Metagene plot showing that hypermethylated m^6^A sites predominantly occur in regions surrounding the stop codon. **(C)** Detection of alternative polyadenylation events using DaPars. The majority of transcripts have a negative ΔPDUI in UCB. **(D)** Detection of alternative polyadenylation events in UCB METTL3 knockout clones. Global reduction of m^6^A leads to lengthening of transcripts. **(E)** Almost all transcripts, for which 3’-UTR shortening was reported, are m^6^A-methylated. **(F)** Comparison of VIRMA mRNA expression between control and UCB tissue samples in our data set. P<0.05, Mann-Whitney-U test. **(G)** Comparison of VIRMA mRNA expression between control (n = 28) and UCB (n = 407) tissue samples. Cancer samples are from the TCGA-BLCA cohort, control samples were combined from the TCGA-BLCA and GTEx cohorts. P<0.05, Mann-Whitney-U test. **(H)** Overview of genetic alterations of VIRMA in the TCGA-BLCA dataset. **(I)** Scatter plot depicting the correlation between the log_2_ copy-number values and VIRMA mRNA expression levels. **(J)** 10-year overall survival analysis of the TCGA-BLCA cohort. Patients were stratified into VIRMA-high (n = 345) and VIRMA-low (n = 65) groups. p=0.0146, log-rank test.

Hypermethylation of m^6^A sites in 3’-UTR and stop codon regions has been associated with VIRMA in HeLa cells (Yue *et al*., 2018). We therefore determined the mRNA expression levels of VIRMA in our sample set. The results showed that VIRMA was significantly (p<0.05, Mann-Whitney-U test) upregulated in UCB (Fig. 5F). Combined TCGA and GTEx cohort analyses further confirmed that VIRMA is significantly (p<0.05, Mann-Whitney-U test) overexpressed in UCB when compared to healthy bladder tissue (Fig. 5G). To explore the underlying mechanism driving VIRMA overexpression, we examined genetic alterations in UCB patients. Notably, VIRMA gene amplification was observed in approximately 6 % of tumors (Fig. 5H), which is consistent with the known amplification rates of other UCB-associated genes, including MYC (2.9-3.3 % (Kluth *et al*, 2023; Zaharieva *et al*, 2005)), FGFR1 (3.7-7 % (Bou Zerdan *et al*, 2023; Helsten *et al*, 2016)), and HER2 (8-8.7 % (Bou Zerdan *et al*., 2023; Fleischmann *et al*, 2011)). The link between VIRMA amplification and overexpression was further supported by a correlation analysis of VIRMA copy number values and mRNA expression levels, which demonstrated a highly significant (p=3.02e-60, Pearson correlation) positive association (Fig. 5I). Finally, survival analysis revealed that UCB patients with higher VIRMA expression had a significantly (p=0.01, log-rank test) worse prognosis (Fig. 5J). These findings suggest that VIRMA can play an important role in UCB progression, through increased expression and hypermethylation of 3’-UTRs.

## Discussion

m^6^A plays a key role in the regulation of transcripts. Altered m^6^A regulation has been linked to oncogenic phenotypes, with numerous studies reporting aberrant m^6^A signatures and dysregulated expression of m^6^A regulators across various cancer types (Barbieri & Kouzarides, 2020; Koch & Lyko, 2024). However, the vast majority of these studies have relied on antibody-dependent methods, which suffer from important limitations. Very recently, initial results from direct RNA-sequencing of m^6^A in clear renal cell carcinoma and MIBC have been interpreted to reflect that altered modification levels may serve as potential biomarkers for patient risk stratification (Li *et al*., 2024; Zhang *et al*., 2024). Direct sequencing can infer the presence and the stoichiometry of m^6^A sites, but it cannot positively determine them in absolute terms. Furthermore, the use of a supervised learning algorithm for m^6^A detection that was trained on RNA samples from plants (Qin *et al*, 2022), and the analysis of modification sites with very low coverage (>2), in combination with low-stringency cutoffs (p<0.1) for differential methylation detection, likely precludes the robust detection and quantification of m^6^A sites (Zhang *et al*., 2024). As such, the findings from these studies can only be considered as preliminary and our understanding of cancer-associated changes in the m^6^A landscape and their consequences appeared greatly limited.

Here, we used GLORI-sequencing of nine UCB tumor samples and nine control samples to establish the first base-resolution maps of the m^6^A in UCB. Sequence motif analyses showed that GLORI detected m^6^A sites predominantly in DRAC(H) consensus sequence motifs, consistent with the known specificity of the m^6^A writer complex (Dominissini *et al*., 2012; Linder *et al*., 2015; Meyer *et al*, 2012). Notably, while cancer-related mutations in the m^6^A writer complex have been suggested to induce m^6^A methylation in non-canonical motifs (Zhang *et al*, 2023), our GLORI analysis of UCB samples showed that m^6^A sites were predominantly detected in the context of the DRAC(H) motif. These findings support earlier studies reporting relatively robust m^6^A profiles across different physiological cell and tissue types (Liu *et al*, 2020; Schwartz *et al*, 2014), and expand them to cancer. However, we also identified systematic alterations between the m^6^A profiles of UCB and control tissues, which allowed their clear and unambiguous separation in dimensionality reduction analyses. A prominent example was the MYC oncogenic transcript, which is known to be regulated in an m^6^A-dependent manner in UCB (Cheng *et al*., 2019), and which we found to be hypermethylated in its CDS and 3’-UTR regions. Furthermore, we found pronounced hypermethylation in transcripts from other cancer genes, such as SMAD3 and BTG2 (Mao *et al*., 2015; Millet & Zhang, 2007; Yuniati *et al*., 2019). These findings suggest the possibility to develop novel biomarkers from the m^6^A profile of UCB, which would address a major unmet clinical need for this tumor entity (Batista *et al*., 2020).

By integrating information about mRNA abundances from standard RNA-sequencing into our GLORI analyses, we determined methylation levels relative to transcript abundance. This novel computational approach allows for a more accurate and comprehensive assessment of the m^6^A epitranscriptomic landscape by avoiding bias from highly abundant or underrepresented transcripts. When comparing the correspondingly adjusted methylation levels of UCB and control tissues, we found a global reduction of m^6^A in UCB, which is consistent with our previous data obtained by LC-MS/MS (Koch *et al*., 2023). Further analyses suggested that this global reduction was primarily caused by the upregulation of abundant, unmethylated transcripts, which diluted the global methylation level, and by the downregulation of abundant, highly methylated transcripts. These observations are consistent with the hypothesis that changes in m^6^A levels across different subcellular compartments, treatments, or tissue samples result from changes in the mRNA metabolism that affect transcript abundance (Shachar *et al*, 2024). Furthermore, our results highlight the limits of standard m^6^A quantification and/or mapping approaches that do not consider transcript abundances.

Simultaneously, our analysis identified a high density of hypermethylated sites in UCB near stop codons, a feature that has been linked to alternative polyadenylation (Yue *et al*., 2018). Alternative polyadenylation is the process by which different isoforms of the same transcript can be formed based on the usage of proximal or distant polyadenylation sites. Thereby, alternative 3’ ends are generated, which affect the fate of the transcript in terms of stability, localization, and translation (Tian & Manley, 2017). Alternative polyadenylation has also been implied in the etiology of cancer (Yuan *et al*, 2021). When analyzing alternative polyadenylation events in our cohort, we found pronounced 3’-UTR shortening of transcripts in UCB, which is consistent with previous findings in UCB and other cancer entities (Xia *et al*, 2014). Conversely, METTL3 knockout and inhibitor-mediated, global depletion of m^6^A in two UCB cell lines resulted in substantial 3’-UTR lengthening, which supports published data showing that methylated transcripts prefer proximal polyadenylation sites (Molinie *et al*, 2016).

Finally, our findings also establish a novel link between the cancer-specific m^6^A epitranscriptomic landscape and the m^6^A writer complex component VIRMA. VIRMA has been reported to be upregulated and amplified across many cancer entities, and its elevated expression has been associated with poor patient prognosis (Destefanis *et al*, 2024; Zhu *et al*, 2021). Furthermore, VIRMA-dependent m^6^A changes have been linked to the progression of certain cancers (Lee *et al*, 2023; Li *et al*, 2023; Yang *et al*, 2024a; Zheng *et al*, 2023; Zhu *et al*, 2024). Our findings are in agreement with these studies and provide important mechanistic support for a functional role of m^6^A in UCB progression. Altogether, our study thus identifies two separate mechanisms that alter the m^6^A epitranscriptomic landscape in cancer (Fig. 6). Compared to earlier antibody-based m^6^A mapping approaches, the highly precise and quantitative results generated by GLORI were critical for uncovering these signatures. Further work will be needed to elucidate the cancer-associated role of VIRMA-driven m⁶A hypermethylation near stop codons in regulating alternative polyadenylation, and to explore the potential of m⁶A profiling for identifying biomarkers for early detection of bladder cancer.

**Figure 6.**
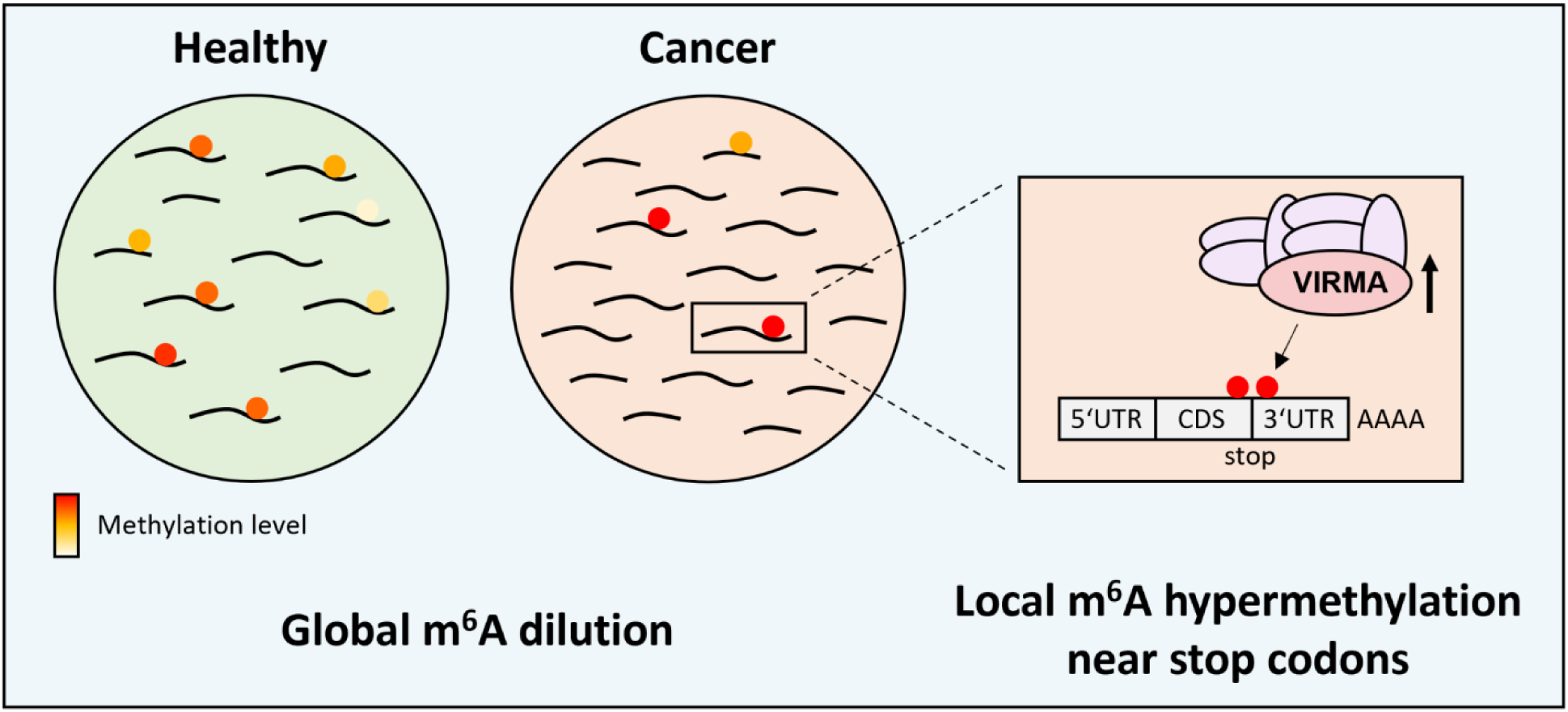
The m^6^A epitranscriptomic landscape of bladder cancer is characterized by global methylation dilution and 3’-UTR hypermethylation. Globally, the upregulation of unmethylated transcripts and the downregulation of highly methylated transcripts resulted in the dilution of m^6^A methylation levels in bladder cancer. Our findings further suggest that the upregulation of VIRMA causes the local hypermethylation of m^6^A modified target transcripts in regions close to the stop codon.

## Methods

### Urothelial carcinoma patients and sample acquisition

This study was performed in adherence to the Declaration of Helsinki. All patients provided informed consent to participate in the molecular characterization of their tissue samples. Additionally, approval of the institutional ethics review board (Ethical Committee II, University of Heidelberg, Germany, reference number: 2015-549N-MA) was taken. Patient information is provided in Table EV 3.

Tumor samples were obtained from transurethral resection of the bladder (TURB) specimens. Non-malignant uroepithelial tissue was taken from cystectomy specimens. Samples were diagnosed by an uropathologist, and tumors were characterized according to the TNM classification for bladder cancer by the Union for International Cancer Control (UICC 2017). Bladder tumors with variant histopathological findings other than urothelial carcinoma were excluded. Patient tissue samples were stored at -20 °C in RNAlater solution until further processing.

### Cell culture

Cell lines were authenticated by single nucleotide polymorphism-profiling, tested for mycoplasma and cultured based on ATCC guidelines. HEK293T and UM-UC-3 cell lines were cultured in DMEM high glucose medium supplemented with 10 % FBS and 1 % P/S. T24 cells were cultured in McCoy’s 5A (modified) medium supplemented with 10 % FBS and 1 % P/S. All cell lines were cultivated as adherent monolayers at 37 °C in a humidified incubator with an atmosphere of 5 % CO_2_.

### Establishment of T24 and UM-UC-3 METTL3 knockout clones

HEK293T cells were transfected with the lentiviral packaging vectors psPAX (Plasmid #12260, Addgene) and pMD2.G (Plasmid #87360, Addgene) as well as the pLentiCRISPR v2 vector (Plasmid #52961, Addgene) using Lipofectamine 2000 (Thermo Fisher Scientific). sgRNAs comprised: Anti-METTL3 sgRNA1: 5’-AGACTAGGATGTCGGACACG-3’. Anti-METTL3 sgRNA2: 5’-CTGGTGGCCCTAAGCCCAGC-3’. Anti-METTL3 sgRNA3: 5’- ATGCTGACCATTCCAAGCTC-3’. Anti-METTL3 sgRNA4: 5’-AAGTGCAAGAATTCTGTGAC-3’. Non-targeting scramble sgRNA: 5’-GCTGACGGCGAGCTTTAGGC-3’. Transfected HEK293T cells were incubated for 48 hours. T24 and UM-UC-3 cells were transduced for 48 hours and further cultivated for the establishment of clonal populations.

### STM2457 treatment of urothelial carcinoma cells

T24 and UM-UC-3 cells were cultured until reaching a confluency of approximately 80 %. The cells were then treated for 48h with DMSO or 50 µM of the METTL3 inhibitor STM2457 (MedChemExpress).

### Isolation and preparation of RNA samples for GLORI-sequencing

RNA isolation and preparation for GLORI were performed as described (Liu *et al*., 2023; Shen *et al*, 2024). Patient tissue samples were homogenized by disruption using the TissueRuptor II (Qiagen). Total RNA from homogenized tissue samples and HEK293T cells was isolated with TRIzol. Small RNA fractions were depleted using the MEGAclear Transcription Clean-Up Kit (Thermo Fisher Scientific). Enrichment of mRNA was performed twice using the Dynabeads mRNA Purification Kit (Thermo Fisher Scientific). mRNA was then fragmented by incubation for 3 min at 94 °C using the NEBNext Magnesium RNA Fragmentation Module (New England Biolabs). The fragmentation step was performed differently from the published version of the protocol which described mRNA fragmentation for 4 min at 94 °C. Fragmented mRNA was then DNase I-treated and purified by the RNA Clean & Concentrator-5 kit (Zymo Research). The concentration as well as the average and peak mRNA fragment lengths were determined by TapeStation (Agilent Technologies).

### Protection, deamination, and deprotection of RNA samples for GLORI-sequencing

All steps were performed as described (Liu *et al*., 2023; Shen *et al*., 2024). For RNA protection, 100 – 200 ng of fragmented mRNA supplemented with a synthetic Spike-In RNA oligonucleotide (Sequence: 5’- AGCGCAGCGAACAUGACACGUGCUCAACAGUGUAGUUGGACUUCCUCGCAUCUAGCCGCUAUGCACGUGCAGCU-3’) were added into protection buffer (1.32 M Glyoxal solution, 50 % DMSO, prepared in water) and incubated for 30 min at 50 °C in a thermal cycler. The RNA was further stabilized by adding 10 µL of freshly prepared saturated boric acid at RT, followed by incubation for 30 min at 50 °C in a thermal cycler. Protected RNA was then added into 50 µL of freshly prepared deamination buffer (1.5 M NaNO2, 80 mM MES (pH 6.0), 1.76 M Glyoxal solution, prepared in water), and incubated for 8 hours at 16 °C in a thermal cycler. After deamination, RNA was purified by ethanol precipitation ON at -80 °C. Pelleted RNA was washed twice using 75 % ethanol and air-dried for 5 min at RT. Then, the RNA pellet was resuspended in 50 µL of deprotection buffer (500 mM triethylammonium acetate (pH 8.6), 47.5 % deionized formamide, prepared in water) and incubated for 10 min at 95 °C in a thermal cycler. Deprotected RNA was purified by ethanol precipitation for at least 30 min at -80 °C. Pelleted RNA was washed twice using 75 % ethanol and air-dried for 5 min at RT. The RNA pellet was then resuspended in 50 µL of water and further purified using the RNA Clean & Concentrator-5 kit (Zymo Research).

### Preparation of GLORI libraries and sequencing

For sequencing library preparation, a two-step RNA end-repair protocol was used. RNA 3’-end dephosphorylation was performed by Antarctic phosphatase (New England Biolabs) treatment in a total reaction volume of 20 µL. The reaction was incubated for 30 min at 37 °C and inactivated for 2 min at 80 °C. Subsequently, RNA 5’-end phosphorylation was performed by T4 Polynucleotide Kinase (New England Biolabs) treatment in a total reaction volume of 50 µL. The reaction was incubated for 30 min at 37 °C, inactivated for 20 min at 65 °C, and purified using the RNA Clean & Concentrator-5 kit (Zymo Research). GLORI-sequencing libraries were then prepared using the NEBNext Small RNA Library Prep Set for Illumina in combination with the NEBNext Multiplex Oligos for Illumina (Index Primer Sets 1 and 3) (New England Biolabs). Size selection of the libraries was performed via TBE-PAGE using 8 % TBE polyacrylamide gels (Thermo Fisher Scientific) selecting sequencing competent molecules in a size range of 160 – 250 bp. Average peak size and concentration of the libraries were determined by TapeStation (Agilent Technologies). Libraries were sequenced on a NovaSeq 6000 platform (Illumina) applying a 100 bp paired-end sequencing protocol. Sequencing was performed by the Next Generation Sequencing Core Facility of the German Cancer Research Center, Heidelberg. Raw sequencing data were then trimmed using Trim Galore (version 0.6.6) and further processed by the GLORI-tools pipeline as described (Liu *et al*., 2023; Shen *et al*., 2024). GLORI-tools is available on GitHub: https://github.com/liucongcas/GLORI-tools. Parameters for the selection of m^6^A sites were set as follows: ≥ 5 variant nucleotides, ≥ 15 coverage of A and G bases, ≥ 0.1 A rate (methylation level > 10 %). All steps of the GLORI-tools pipeline were executed using the following software: python (version 3.10.1), samtools (version 1.19), STAR (version 2.7.10a), bowtie (version 1.3.0). The human genome (GRCh38) and transcriptome (GCF_000001405.39) reference files were downloaded from UCSC.

### Motif analysis

The coordinates and flanking sequences of GLORI-derived m^6^A sites were extracted using bedtools getfasta (version 2.25.0) and the human genome reference file (GRCh38). Sequence motifs were then identified using Hypergeometric Optimization of Motif EnRichment (HOMER, version 4.11). For visualization, the motif representations were converted into position weight matrices using DiffLogo (version 2.20.0).

### Differential methylation analysis

The R package methylSig (version 1.7.0) was used to identify differentially methylated sites comparing control and cancer tissues. For the analysis, an absolute methylation level cutoff of 10 % and a p-value cutoff of 0.05 were selected. The statistical significance was determined using beta binomial models. Gene ontology analyses were performed using ShinyGO (version 0.81, (Ge *et al*, 2020)).

### Bulk RNA-sequencing

Total RNA from tissue samples and cell lines was isolated using TRIzol. To remove residual gDNA contaminants, total RNA samples were DNase I-digested and further purified using the RNA Clean & Concentrator-5 kit (Zymo Research). Library preparation and sequencing were performed by the Next Generation Sequencing Core Facility of the German Cancer Research Center, Heidelberg. The samples were sequenced on a NovaSeq 6000 platform (Illumina) applying a 100 bp paired-end sequencing protocol. Reads were trimmed using Trim Galore (version 0.6.6) and mapped to the human reference genome (GRC38) using HISAT2 (version 2.2.1).

### Differential gene expression analysis

Aligned reads from bulk RNA-sequencing experiments were counted by the featureCounts function (2.0.6), normalized, and differential gene expression analysis performed via DESeq2 (version 1.42.0).

### Analysis of alternative polyadenylation events

For the de novo identification of alternative polyadenylation sites, aligned reads from bulk RNA-sequencing experiments were analysed using the DaPars algorithm, version 1.0.0 (Xia *et al*., 2014). For the analyses, a coverage cutoff of 30, an FDR cutoff of 0.05, an absolute PDUI difference cutoff of 0.1, and an absolute fold change cutoff of 0.59 were selected.

### Weighted global methylation level analysis

For the determination of the weighted global methylation level in the individual samples, m^6^A methylation data from GLORI-sequencing and transcript expression data from RNA-sequencing were integrated. The methylation level of each transcript was aggregated and then multiplied by the TPM-normalized expression level of the same transcript. These individual, weighted methylation levels were then summed up to compute the weighted global methylation level.

### Databank VIRMA expression analysis

VIRMA mRNA expression data were downloaded from the UCSC database for the combined The Cancer Genome Atlas bladder urothelial carcinoma (TCGA-BLCA) cohort and the Genotype-Tissue Expression (GTEx) project (https://xenabrowser.net/). Expression data were DESeq2-normalized and log_2_-transformed (log_2_(value + 1)). Mann-Whitney-U test was assessed to test for statistical significance between control and UCB tissues.

### Association analysis of VIRMA genomic alterations and expression

Genetic alteration data and the corresponding mRNA expression data for VIRMA were downloaded from the cBioPortal database (https://www.cbioportal.org/) for TCGA-BLCA cohort. VIRMA mRNA expression values were preprocessed from RNA-sequencing by Expectation-Maximization (RSEM) data generated by the TCGA RNASeqV2 pipeline (Illumina HiSeq), which were batch-normalized and log_2_-transformed (log_2_(value + 1)). Different copy-number alterations (shallow deletion, diploid, gain, amplification) were identified based on the Genomic Identification of Significant Targets in Cancer (GISTIC) method. Pearson correlation analysis was performed to assess the association between log_2_ copy-number values and VIRMA mRNA expression levels.

### Survival analysis

Kaplan-Meier overall survival analysis was performed using the lifelines python package. Patients were stratified into VIRMA-high and VIRMA-low mRNA expression groups using an optimal cutoff. Statistical significance between two groups was assessed using the log-rank test.

## Data availability

Data generated in this study have been deposited in the GEO database under the accession numbers GSE281749 and GSE281750.

## Acknowledgements

We thank the Genomics and Proteomics Core Facility of the German Cancer Research Center, particularly Franziska Petermann and Panagiotis Provataris for their support. We also thank Sandra Blanco for critically reading the manuscript. This work was supported by a grant from Deutsche Forschungsgemeinschaft (DFG, German Research Foundation) to F.L. (project number 439669440 TRR319 RMaP TP A01). Additional funding was provided by the DKFZ-Hector Seed Funding Program (project UCGLORI).

## Author contributions

J.K. Data curation, Formal Analysis, Investigation, Methodology, Software, Validation, Visualization, Writing – original draft; J.X. Formal Analysis, Investigation, Visualization; F.B. Formal Analysis, Investigation, Software, Validation, Visualization; V.C.C. Supervision; Ma.Ne. Funding acquisition, Resources, Writing – review & editing; K.N. Resources; Ma.Ni. Conceptualization, Funding acquisition, Resources, Writing – review & editing; P.E. Resources, Writing – review & editing; M.S.M. Resources; M.R.P. Funding acquisition, Supervision, Writing – review & editing; F.L. Conceptualization, Data curation, Funding acquisition, Project administration, Resources, Supervision, Writing – original draft.

## References

Barbieri I, Kouzarides T (2020) Role of RNA modifications in cancer. Nat Rev Cancer 20: 303–322

Batista R, Vinagre N, Meireles S, Vinagre J, Prazeres H, Leão R, Máximo V, Soares P (2020) Biomarkers for Bladder Cancer Diagnosis and Surveillance: A Comprehensive Review. Diagnostics (Basel) 10

Berdik C (2017) Unlocking bladder cancer. Nature 551: S34–s35

Bou Zerdan M, Bratslavsky G, Jacob J, Ross J, Huang R, Basnet A (2023) Urothelial Bladder Cancer: Genomic Alterations in Fibroblast Growth Factor Receptor. Mol Diagn Ther 27: 475–485

Boulias K, Greer EL (2023) Biological roles of adenine methylation in RNA. Nature Reviews Genetics 24: 143–160

Cheng M, Sheng L, Gao Q, Xiong Q, Zhang H, Wu M, Liang Y, Zhu F, Zhang Y, Zhang X et al (2019) The m(6)A methyltransferase METTL3 promotes bladder cancer progression via AFF4/NF-κB/MYC signaling network. Oncogene 38: 3667–3680

Deng L-J, Deng W-Q, Fan S-R, Chen M-F, Qi M, Lyu W-Y, Qi Q, Tiwari AK, Chen J-X, Zhang D-M et al (2022) m6A modification: recent advances, anticancer targeted drug discovery and beyond. Mol Cancer 21: 52

Deng X, Qing Y, Horne D, Huang H, Chen J (2023) The roles and implications of RNA m(6)A modification in cancer. Nat Rev Clin Oncol 20: 507–526

Destefanis E, Sighel D, Dalfovo D, Gilmozzi R, Broso F, Cappannini A, Bujnicki JM, Romanel A, Dassi E, Quattrone A (2024) The three YTHDF paralogs and VIRMA are the major tumor drivers among the m^6^A core genes in a pan-cancer analysis. bioRxiv: 2024.2006.2013.598899

Dominissini D, Moshitch-Moshkovitz S, Schwartz S, Salmon-Divon M, Ungar L, Osenberg S, Cesarkas K, Jacob-Hirsch J, Amariglio N, Kupiec M et al (2012) Topology of the human and mouse m6A RNA methylomes revealed by m6A-seq. Nature 485: 201–206

Dyrskjøt L, Hansel DE, Efstathiou JA, Knowles MA, Galsky MD, Teoh J, Theodorescu D (2023) Bladder cancer. Nat Rev Dis Primers 9: 58

Flamand MN, Tegowski M, Meyer KD (2023) The Proteins of mRNA Modification: Writers, Readers, and Erasers. Annu Rev Biochem 92: 145–173

Fleischmann A, Rotzer D, Seiler R, Studer UE, Thalmann GN (2011) Her2 amplification is significantly more frequent in lymph node metastases from urothelial bladder cancer than in the primary tumours. Eur Urol 60: 350–357

Ge SX, Jung D, Yao R (2020) ShinyGO: a graphical gene-set enrichment tool for animals and plants. Bioinformatics 36: 2628–2629

Gill E, Perks CM (2024) Mini-Review: Current Bladder Cancer Treatment-The Need for Improvement. Int J Mol Sci 25

Goriki A, Seiler R, Wyatt AW, Contreras-Sanz A, Bhat A, Matsubara A, Hayashi T, Black PC (2018) Unravelling disparate roles of NOTCH in bladder cancer. Nat Rev Urol 15: 345–357

He PC, He C (2021) m(6) A RNA methylation: from mechanisms to therapeutic potential. Embo j 40: e105977

Helm M, Lyko F, Motorin Y (2019) Limited antibody specificity compromises epitranscriptomic analyses. Nat Commun 10: 5669

Helsten T, Elkin S, Arthur E, Tomson BN, Carter J, Kurzrock R (2016) The FGFR Landscape in Cancer: Analysis of 4,853 Tumors by Next-Generation Sequencing. Clin Cancer Res 22: 259–267

Hewel C, Wierczeiko A, Miedema J, Hofmann F, Weißbach S, Dietrich V, Friedrich J, Holthöfer L, Haug V, Mündnich S et al (2025) Direct RNA sequencing enables improved transcriptome assessment and tracking of RNA modifications for medical applications. bioRxiv: 2024.2007.2025.605188

Kluth M, Hitzschke M, Lennartz M, Blessin NC, Sauter G, Plage H, Klatte T, Schlomm T, Marx AH, Rink M et al (2023) MYC amplifications are a common event in urothelial bladder carcinomas associated with an aggressive tumor phenotype. American Journal of Clinical Pathology 160: S91–S92

Knowles MA, Hurst CD (2015) Molecular biology of bladder cancer: new insights into pathogenesis and clinical diversity. Nat Rev Cancer 15: 25–41

Koch J, Lyko F (2024) Refining the role of N6-methyladenosine in cancer. Current Opinion in Genetics & Development 88: 102242

Koch J, Neuberger M, Schmidt-Dengler M, Xu J, Carneiro VC, Ellinger J, Kriegmair MC, Nuhn P, Erben P, Michel MS et al (2023) Reinvestigating the clinical relevance of the m(6)A writer METTL3 in urothelial carcinoma of the bladder. iScience 26: 107300

Lan Q, Liu PY, Haase J, Bell JL, Hüttelmaier S, Liu T (2019) The Critical Role of RNA m(6)A Methylation in Cancer. Cancer Res 79: 1285–1292

Lee Q, Song R, Phan DAV, Pinello N, Tieng J, Su A, Halstead JM, Wong ACH, van Geldermalsen M, Lee BS et al (2023) Overexpression of VIRMA confers vulnerability to breast cancers via the m(6)A-dependent regulation of unfolded protein response. Cell Mol Life Sci 80: 157

Li H, Li C, Zhang Y, Jiang W, Zhang F, Tang X, Sun G, Xu S, Dong X, Shou J et al (2024) Comprehensive analysis of m(6) A methylome and transcriptome by Nanopore sequencing in clear cell renal carcinoma. Mol Carcinog 63: 677–687

Li N, Zhu Z, Deng Y, Tang R, Hui H, Kang Y, Rana TM (2023) KIAA1429/VIRMA promotes breast cancer progression by m(6) A-dependent cytosolic HAS2 stabilization. EMBO Rep 24: e55506

Linder B, Grozhik AV, Olarerin-George AO, Meydan C, Mason CE, Jaffrey SR (2015) Single-nucleotide-resolution mapping of m6A and m6Am throughout the transcriptome. Nature Methods 12: 767–772

Lister R, Ecker JR (2009) Finding the fifth base: genome-wide sequencing of cytosine methylation. Genome Res 19: 959–966

Liu C, Sun H, Yi Y, Shen W, Li K, Xiao Y, Li F, Li Y, Hou Y, Lu B et al (2023) Absolute quantification of single-base m6A methylation in the mammalian transcriptome using GLORI. Nature Biotechnology 41: 355–366

Liu J, Li K, Cai J, Zhang M, Zhang X, Xiong X, Meng H, Xu X, Huang Z, Peng J et al (2020) Landscape and Regulation of m(6)A and m(6)Am Methylome across Human and Mouse Tissues. Mol Cell 77: 426–440.e426

Lopez-Beltran A, Cookson MS, Guercio BJ, Cheng L (2024) Advances in diagnosis and treatment of bladder cancer. Bmj 384: e076743

Mao B, Zhang Z, Wang G (2015) BTG2: a rising star of tumor suppressors (review). Int J Oncol 46: 459–464

McIntyre ABR, Gokhale NS, Cerchietti L, Jaffrey SR, Horner SM, Mason CE (2020) Limits in the detection of m6A changes using MeRIP/m6A-seq. Scientific Reports 10: 6590

Meyer KD, Saletore Y, Zumbo P, Elemento O, Mason CE, Jaffrey SR (2012) Comprehensive analysis of mRNA methylation reveals enrichment in 3’ UTRs and near stop codons. Cell 149: 1635–1646

Millet C, Zhang YE (2007) Roles of Smad3 in TGF-beta signaling during carcinogenesis. Crit Rev Eukaryot Gene Expr 17: 281–293

Molinie B, Wang J, Lim KS, Hillebrand R, Lu ZX, Van Wittenberghe N, Howard BD, Daneshvar K, Mullen AC, Dedon P et al (2016) m(6)A-LAIC-seq reveals the census and complexity of the m(6)A epitranscriptome. Nat Methods 13: 692–698

Qin H, Ou L, Gao J, Chen L, Wang JW, Hao P, Li X (2022) DENA: training an authentic neural network model using Nanopore sequencing data of Arabidopsis transcripts for detection and quantification of N(6)-methyladenosine on RNA. Genome Biol 23: 25

Richters A, Aben KKH, Kiemeney L (2020) The global burden of urinary bladder cancer: an update. World J Urol 38: 1895–1904

Saginala K, Barsouk A, Aluru JS, Rawla P, Padala SA, Barsouk A (2020) Epidemiology of Bladder Cancer. Med Sci (Basel) 8

Schwartz S, Mumbach MR, Jovanovic M, Wang T, Maciag K, Bushkin GG, Mertins P, Ter-Ovanesyan D, Habib N, Cacchiarelli D et al (2014) Perturbation of m6A writers reveals two distinct classes of mRNA methylation at internal and 5’ sites. Cell Rep 8: 284–296

Shachar R, Dierks D, Garcia-Campos MA, Uzonyi A, Toth U, Rossmanith W, Schwartz S (2024) Dissecting the sequence and structural determinants guiding m6A deposition and evolution via inter- and intra-species hybrids. Genome Biol 25: 48

Shen W, Sun H, Liu C, Yi Y, Hou Y, Xiao Y, Hu Y, Lu B, Peng J, Wang J et al (2024) GLORI for absolute quantification of transcriptome-wide m6A at single-base resolution. Nature Protocols 19: 1252–1287

Siegel RL, Giaquinto AN, Jemal A (2024) Cancer statistics, 2024. CA Cancer J Clin 74: 12–49

Sui X, Lei L, Chen L, Xie T, Li X (2017) Inflammatory microenvironment in the initiation and progression of bladder cancer. Oncotarget 8: 93279–93294

Sun H, Lu B, Zhang Z, Xiao Y, Zhou Z, Xi L, Li Z, Jiang Z, Zhang J, Wang M et al (2025) Mild and ultrafast GLORI enables absolute quantification of m6A methylome from low-input samples. Nature Methods

Sylvester RJ, van der Meijden AP, Oosterlinck W, Witjes JA, Bouffioux C, Denis L, Newling DW, Kurth K (2006) Predicting recurrence and progression in individual patients with stage Ta T1 bladder cancer using EORTC risk tables: a combined analysis of 2596 patients from seven EORTC trials. Eur Urol 49: 466–465; discussion 475-467

Tian B, Manley JL (2017) Alternative polyadenylation of mRNA precursors. Nat Rev Mol Cell Biol 18: 18–30

Xia Z, Donehower LA, Cooper TA, Neilson JR, Wheeler DA, Wagner EJ, Li W (2014) Dynamic analyses of alternative polyadenylation from RNA-seq reveal a 3’-UTR landscape across seven tumour types. Nat Commun 5: 5274

Yang K, Zhong Z, Zou J, Liao JY, Chen S, Zhou S, Zhao Y, Li J, Yin D, Huang K et al (2024a) Glycolysis and tumor progression promoted by the m(6)A writer VIRMA via m(6)A-dependent upregulation of STRA6 in pancreatic ductal adenocarcinoma. Cancer Lett 590: 216840

Yang L, Ying J, Tao Q, Zhang Q (2024b) RNA N(6)-methyladenosine modifications in urological cancers: from mechanism to application. Nat Rev Urol 21: 460–476

Yuan F, Hankey W, Wagner EJ, Li W, Wang Q (2021) Alternative polyadenylation of mRNA and its role in cancer. Genes Dis 8: 61–72

Yue Y, Liu J, Cui X, Cao J, Luo G, Zhang Z, Cheng T, Gao M, Shu X, Ma H et al (2018) VIRMA mediates preferential m(6)A mRNA methylation in 3’UTR and near stop codon and associates with alternative polyadenylation. Cell Discov 4: 10

Yuniati L, Scheijen B, van der Meer LT, van Leeuwen FN (2019) Tumor suppressors BTG1 and BTG2: Beyond growth control. J Cell Physiol 234: 5379–5389

Zaccara S, Ries RJ, Jaffrey SR (2019) Reading, writing and erasing mRNA methylation. Nature Reviews Molecular Cell Biology 20: 608–624

Zaharieva B, Simon R, Ruiz C, Oeggerli M, Mihatsch MJ, Gasser T, Sauter G, Toncheva D (2005) High-throughput tissue microarray analysis of CMYC amplificationin urinary bladder cancer. Int J Cancer 117: 952–956

Zhang C, Tunes L, Hsieh MH, Wang P, Kumar A, Khadgi BB, Yang Y, Doxtader KA, Herrell E, Koczy O et al (2023) Cancer mutations rewire the RNA methylation specificity of METTL3-METTL14. bioRxiv

Zhang L, Chen Z, Sun G, Li C, Wu P, Xu W, Zhu H, Zhang Z, Tang Y, Li Y et al (2024) Dynamic landscape of m6A modifications and related post-transcriptional events in muscle-invasive bladder cancer. J Transl Med 22: 912

Zheng ZQ, Huang ZH, Liang YL, Zheng WH, Xu C, Li ZX, Liu N, Yang PY, Li YQ, Ma J et al (2023) VIRMA promotes nasopharyngeal carcinoma, tumorigenesis, and metastasis by upregulation of E2F7 in an m6A-dependent manner. J Biol Chem 299: 104677

Zhu C, Cheng Y, Yu Y, Zhang Y, Ren G (2024) VIRMA promotes the progression of head and neck squamous cell carcinoma by regulating UBR5 mRNA and m6A levels. Biomol Biomed 24: 1244–1257

Zhu W, Wang JZ, Wei JF, Lu C (2021) Role of m6A methyltransferase component VIRMA in multiple human cancers (Review). Cancer Cell Int 21: 172

